# A global high-density chromatin interaction network reveals functional long-range and trans-chromosomal relationships

**DOI:** 10.1101/2022.03.24.485503

**Authors:** Ruchi Lohia, Nathan Fox, Jesse Gillis

## Abstract

Chromatin contacts are essential for gene-expression regulation, however, obtaining a high-resolution genome-wide chromatin contact map is still prohibitively expensive owing to large genome sizes and the quadratic scale of pairwise data. Chromosome conformation capture (3C) based methods such as Hi-C have been extensively used to obtain chromatin contacts. However, since the sparsity of these maps increases with an increase in genomic distance between contacts, long-range or trans chromatin contacts are especially challenging to sample.

Here, we created a high density reference genome-wide chromatin contact map using a meta-analytic approach. We integrate 3600 Human, 6700 Mouse, and 500 Fly 3C experiments to create species-specific meta-3C contact maps with 304 billion, 193 billion, and 19 billion contacts in respective species. We validate that meta-3C are uniquely powered to capture functional chromatin contacts in both cis and trans. Unlike individual experiments, meta-3C gene contacts predict gene coexpression for long-range and trans chromatin contacts. Similarly, for long-range cis-regulatory interactions, meta-3C contacts outperform all individual experiments, providing an improvement over the conventionally used linear genomic distance-based association. Assessing between species, we find patterns of chromatin contacts conservation in both cis and trans and strong associations with coexpression even in species for which 3C data is lacking.

We have generated an integrated chromatin interaction network which complements a large number of methodological and analytic approaches focused on improved specificity or interpretation. This high-depth “super-experiment” is surprisingly powerful in capturing long-range functional relationships of chromatin interactions, which are now able to predict coexpression, expression quantitative trait loci (eQTL), and cross-species relationships.

## Introduction

The physical associations generated by chromatin contacts are a critical factor to regulate and determine gene-expression patterns (1–3). Functional chromatin contacts can form across a wide range of genomic distances within a chromosome (cis) or across a chromosome (trans). Although trans contacts are non-random (4) and there is evidence of trans-regulatory interactions (5, 6), studying the functional role of these interactions is difficult due to the high sparsity of available contact maps in trans.

Obtaining high-density contact maps at all genomic distances and in trans is not yet feasible with most existing maps being essentially probabilistic in nature, capturing some fraction of likely-present contacts in a distance-dependent manner. Genome-wide contact maps can be obtained using chromosome conformation capture (3C)-based technologies such as Hi-C (7). However, due to large genome sizes and the quadratic scale of pairwise data, obtaining these maps at high resolution would require prohibitively expensive sequencing at even 1X depth in the pairwise space. Capturing long-range and trans chromatin contacts is made more difficult since the frequency of contacts decreases with an increase in genomic distance between contacting loci in cis (7). And in trans, the contacts are at least 2 orders of magnitude less frequent (4) while also having a larger search space than cis.

To overcome the sequencing-depth barrier targeted 3C-based techniques such as ChiA-PET (8) and Capture-Hi-C (9) are widely used to obtain high-resolution contacts maps for specific proteins or selected loci respectively. Alternatively, several in-silico methods have taken the advantage of existing limited resolution contact maps to either generate higher resolution maps using machine learning approaches (10–13) and/or detect statistically significant interactions by background fitting (14, 15). However, with a few exceptions (16, 17), most of the available methods are only tested to enhance cis interactions because longer range interactions are essentially unavailable within any given data set.

In this work, we propose a meta-analysis approach where we leverage several hundreds of available CC-maps generated from 3C-based experiments to create a dense genome-wide CC-map for three species; Human, Mouse, and Fly We show that these maps are valuable for capturing long-range and trans-chromosomal interactions. We evaluated the effectiveness of contact maps using three criteria; CC-maps were used to predict 1] gene-expression profiles, 2] target genes for eQTLs, and 3] conservation across pairs of species (Human-Mouse, Human-Fly, and Mouse-Fly). Our reference networks complement a very diverse array of efforts in genomics, from those focused on more targeted experiments in 3C which now have an overall “null” with which to compare individual results, to genome interpretation methods, whether interpreting variants, expression patterning, or regulatory sequence.

## Results

### Meta-3C network predicts coexpression at greater resolution and scale than individual networks

In brief, for building the meta-3C network, we uniformly processed 3619, 6732, and 487 3C runs for Human, Mouse, and Fly respectively (Figure 1C). The runs were obtained after querying Sequence Read Archive (SRA) with field limitations of given species and Hi-C as experiment strategy. A genome-wide interaction matrix was created for each run after mapping the reads to the same reference genome for each species. Within SRA, all the runs (SRR) belonging to a study are grouped together as project (SRP). A project can consist of multiple runs, which can include biological or technical replicates across multiple tissues or cell types. All the interaction matrices within a project were aggregated to create a project-level aggregate. There were 119, 33, and 29 projects for Human, Mouse, and Fly respectively. The meta-3C map was created after further aggregating all processed runs within their respective species (Figure 1B). For subsequent analysis, the genome-wide contact matrix was mapped to genes (see Methods) to create networks where nodes are genes and edges are the interaction frequency between genes. The genome-wide networks were divided into cis and trans depending on if the edge connects two genes in the same chromosome or different chromosome respectively (Figure 1A). To validate the predictive power of the meta-3C network, we benchmarked it against networks inferred from individual projects for each species.

**Fig. 1.**
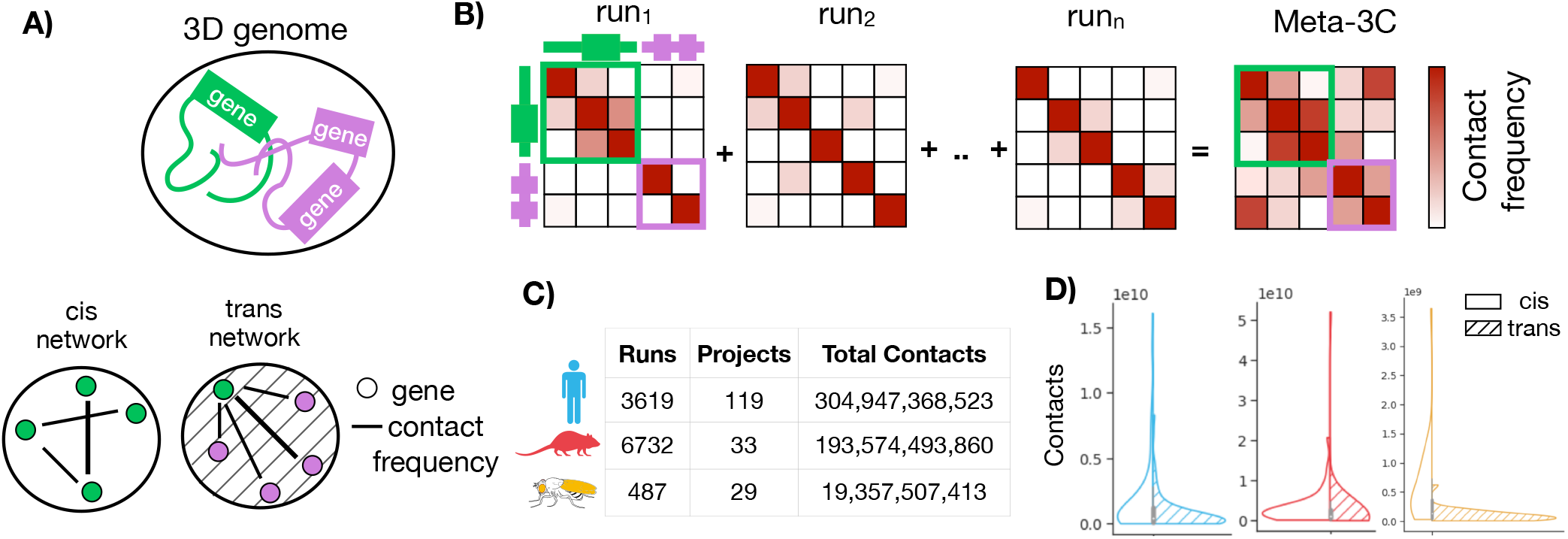
Creating meta-3C network. A) Genes are co-localized in chromatin 3D structure through frequent chromatin contacts. The structure can be represented with networks where nodes are genes and edges are interaction frequencies. The cis and trans networks consist of intra-chromosome and inter-chromosome edges respectively. B) Individual 3C experiments are aggregated to create a meta-3C network. C) Runs, projects, and contacts in Human, Mouse, and Fly meta-3C networks. D) Total contacts (sequencing depth) distribution across projects in cis and trans for each species.

As our first performance test, we assessed the tendency for spatially co-localized to be co-expressed (18, 19), using previously derived shared patterns of expression in independent data (20). The underlying hypothesis is that spatial proximity may be a useful way to organize regulatory relationships, as in the case of linear sequence, thus yielding shared spatial relationships for genes that are co-expressed. Thus, while perfect performance at predicting coexpression is not expected, the genome-wide scale of the assessment makes it useful for assessing cis and trans effects. For each gene, we measure the ability of interaction frequency to predict the gene’s top 1% coexpression partners (Figure 2A). We call this measure “contact coexpression” and is expressed as an AUC (Area Under the ROC Curve) with possible values ranging between 0 and 1. A score of 1 indicates that interaction frequency perfectly predicts coexpression; 0.5 indicates no relationship. We evaluated the contact coexpression as a function of the sequencing depth of the 3C network (Figure 2B and Figure 2C). We find that performance is linearly dependent on the log of sequencing depth and meta-analysis provides additional coverage. We find that in cis the best powered individual experiments are close to the saturation depth that maximizes performance (Figure 2B), although performing substantially worse in trans. In trans, the meta-3C network acts like a “super-experiment”, where the additional coverage fully converts into substantial additional performance (Figure 2C). We found similar results for Mouse (Figure S1) and Fly (Figure S2).

**Fig. 2.**
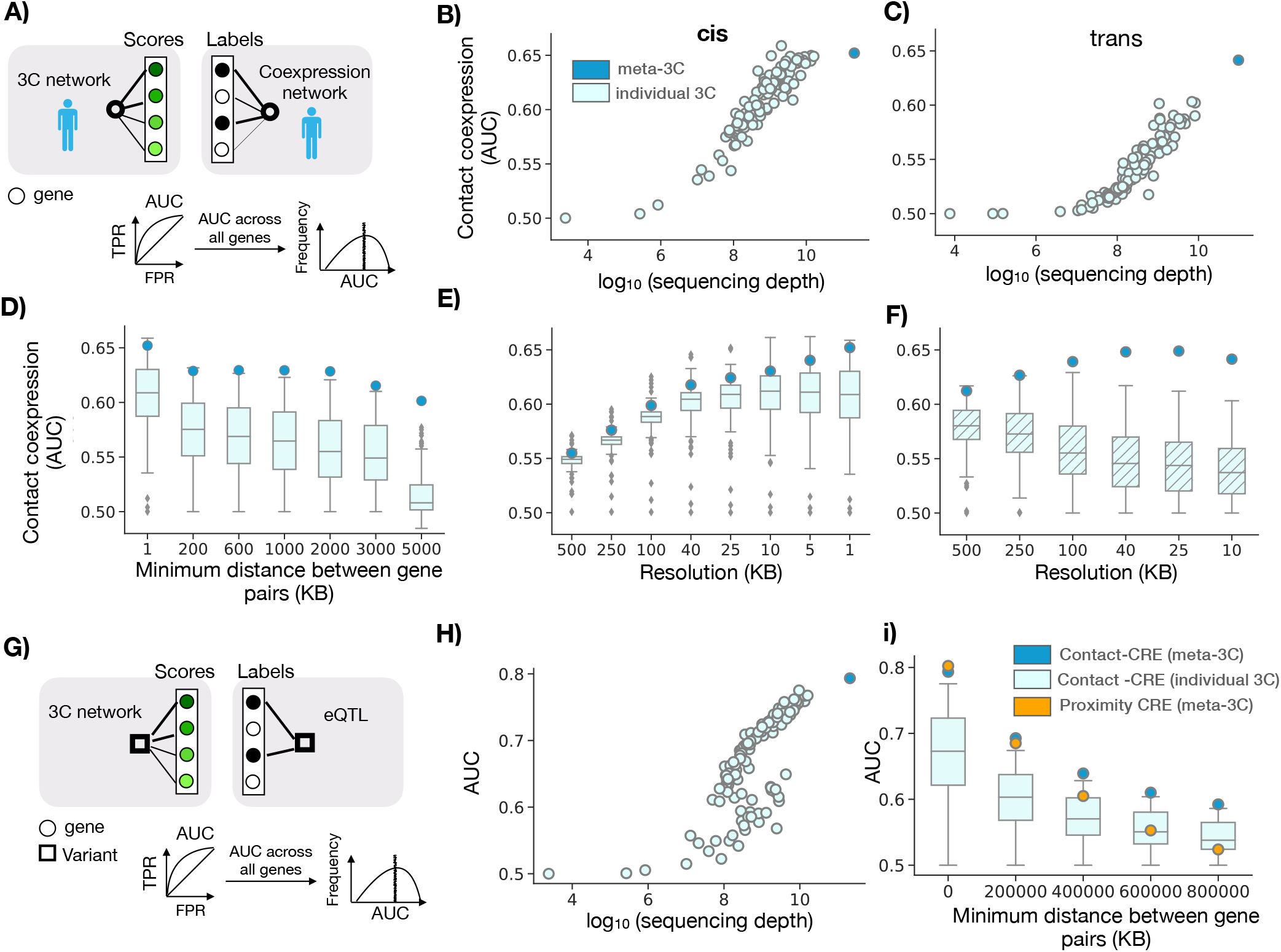
Meta-3C network benchmarking. A) Contact coexpression metric schematic. Circles represent genes and lines represent edges of that gene in respective networks. For each target gene, we use its ranked edges in 3C network to predict the top 1% of its edges in the coexpression network. We perform this task for every gene and then report the average across all genes. B) The circles are contact coexpression for individual and meta3C network in cis as a function of sequencing depth at 1KB resolution C) Same as B) but in trans and at 10KB resolution. D) The boxplot shows the distribution of contact coexpression in cis for each project at various distance thresholds. E) The boxplot shows the distribution of contact coexpression for each project at various resolutions in cis. Circles represent the performance of cis meta-3C network. F) Same as E) but using trans networks. G) Contact-CRE and proximity CRE metric schematic. For contact-CRE and proximity-CRE, for each variant, the edges are ranked by contact frequency or inverse of the genomic distance from the variant respectively. The labels are obtained from eQTL associations (Methods). We perform this task for every variant and then report the average across all variants. H) Contact-CRE for individual project and meta-3C network i) Contact-CRE and Proximity-CRE for meta-3C network, distribution across individual networks at various minimum distance thresholds.

In order to validate that the meta-3C network has more uniform coverage, we compared the contact coexpression of individual 3C networks and meta-3C networks at various linear distance thresholds in cis. We find that for long-range contacts meta-3C network performs better than every individual 3C network (Figure 2D). For both individual networks and meta-3C network, the performance decreases in the absence of short-range contacts. This could be due to a higher number of short-range regulatory interactions or due to the similarity of the chromatin environment for nearby genes. The contact coexpression is dependent on the resolution of the 3C network used and therefore we compared the performance of individual 3C and meta-3C networks at various resolutions. We find that for the individual networks performance increases with an increase in resolution, plateaus, and then slightly falls off in cis (Figure 2D). In essence, improved resolution is useful in cis because the coverage is adequate for it to provide useful signal until the very finest resolution where most experiments begin to decline, although the meta-3C network continues to slightly increase, as might be expected. In contrast, in trans (Figure 2E) the performance monotonically falls with an increase in resolution for individual experiments. However, the in trans pattern for meta-3C networks strongly resembles that of individual experiments in cis, increasing and then plateauing with improvements in resolution (Figure 2D, Figure 2E). This suggests unlike individual networks, meta-3C networks are dense enough to be analyzed at high resolutions even in trans. We found similar results for Mouse (Figure S1) and Fly (Figure S2).

At a large genomic scale, the genome is spatially segregated into two compartments; A and B (7). These compartments can be identified using long-range chromatin contacts. Since the interaction between genes is largely constrained to occur within the same compartment we asked the question if the contact coexpression performance of trans meta-3C network can be explained with A/B compartments. In each individual network, we identified the compartment of each gene and then binarized the compartment preference for each gene pair: gene pair found in the same compartment or different compartment. For each gene, we measure the ability of same compartment frequency to predict the gene’s top 1% coexpression partners (Figure S3). We call this measure “compartment coexpression” and is expressed as an AUC with possible values ranging between 0 and 1. A score of 1 indicates that same compartment frequency perfectly predicts coexpression; 0.5 indicates no relationship. We compared this performance with contact coexpression performance (Figure S3). We find that compartment co-expression performance is a small fraction of contact co-expression, suggesting either more complex sub-compartmental relationships or other trans interactions contribute.

### Meta-3C network effectively capture more eQTL interactions

For our second performance assessment, we tested the hypothesis that genetic variants (eVariant) regulate gene expression of the target gene (eGene) via physical contact (21, 22). The set of eQTLs was obtained from GTEx (Methods). For each eVariant, the interaction frequency with all genes falling in unique contact map bins at 1KB resolution was used to predict the eGene (Figure 2G). This is termed “contact-CRE’ where CRE stands for cis-regulatory elements and is expressed as an AUC with possible values ranging between 0-1, with 1 and 0 meaning that the eVariant and target eGenes have the highest and lowest interaction frequency respectively when compared to all eVariant and non-eGenes interactions. Similar to the previous benchmarking test we find that performance is linearly dependent on the log of sequencing depth and meta-analysis provides additional coverage; meta-3C network has higher performance when compared to any of the individual networks (Figure 2H). This emphasizes the significance of dense contact networks in identifying regulatory interactions.

We further evaluated the ability of meta-3C networks to predict target genes for variants by comparing their performance with a linear genomic distance-based predictor, the current standard approach. The distance between the variant and gene transcription start site (TSS) remains almost the only metric widely used to annotate target genes for variants (23). For each eVariant the inverse of linear distance (1/TSS) with all genes is ranked and then used to predict the eGene (Figure 2G). This is termed “proximity-CRE’ and is expressed as an AUC with possible values ranging between 0-1, with 1 and 0 meaning that eGenes are the closest and farthest from the eVariant respectively. We compared contact-CRE of individual 3C networks, and meta-3C networks at various linear distance thresholds (Figure 2I). We reassuringly find that meta-3C network outperforms individual networks at all distance thresholds. Furthermore, the performance for both contact-CRE and proximity-CRE decreases in the absence of short-range contacts. This is in agreement with our previous observation where we find that contact coexpression decreases in the absence of short-range contacts.

### Trans-chromosomal chromatin contacts show evolutionary conservation

Having established that meta-3C networks are well powered to capture meaningful contacts, we now use them to study the conservation of genomic contacts between species. Since chromatin contacts regulate gene expression, it is reasonable to expect some conservation of these contacts across species even in the context of large scale genomic alteration and, in the reverse, divergence in contacts across species can help explain regulatory evolution (24, 25). We evaluated the conservation of contacts across species in three different ways; we compare the contact coexpression scores for ortholog genes in each species pair, we use the 3C network of one species to predict either 3C network (“contact conservation”) or coexpression network in another species.

Before directly comparing the contact map across species we first compared the contact coexpression scores for 1:1 orthologous genes across species. We find a strong linear relationship between Human and Mouse scores and a somewhat weaker relationship between Human and Fly scores in both cis (Figure S4A) and trans (Figure 3A). This suggests that if a gene is spatially co-regulated in one species, it is likely to be spatially co-regulated across other species.

**Fig. 3.**
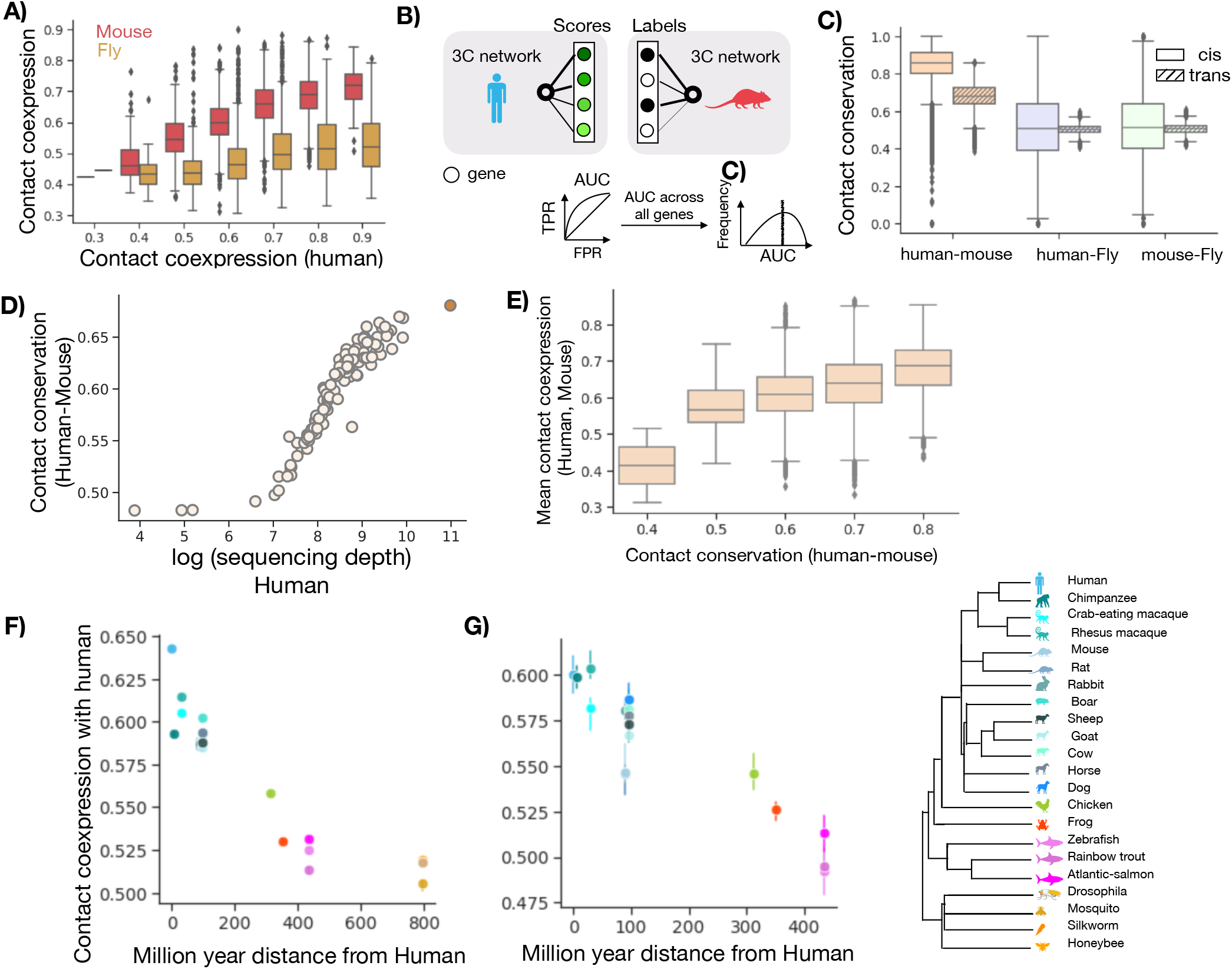
Chromatin contacts are conserved across species in both cis and trans. A) Contact coexpression in trans for 1:1 orthologs in Human-Mouse and Human-Fly. B) Contact conservation schematic. For each gene ranked edges in Human 3C network are used to predict the top 10% of Mouse 3C network edges. This task is repeated for each gene and in both directions and we report the average AUC. C) The distribution of contact conservation score using meta-3C network for various pairs of species. D) Human-Mouse contact conservation for various projects and meta-3C network as a function of sequencing depth. E) Avg contact coexpression score for each gene in Human and Mouse as a function of contact conservation. F) The median performance across all genes when the Human meta-3C network is used to predict coexpression across other species. G) Same as F) but only using the same set of ortholog genes across species. The error bars represent a 68% confidence interval.

We next characterized the degree to which gene contacts are conserved by directly comparing the meta-3C network across species (Figure 3B). Each gene’s shared neighborhood is defined by ranking all edges in the chromatin contact network and then using it to predict the gene’s top 10% of edges in another species. We call this “contact conservation” and again, treat it as a prediction task with 1 meaning perfect contact conservation, 0.5 consistent with random reordering of neighborhoods, and 0 meaning that contacting partners have reversed. For trans conservation score, only the trans gene pairs in both species are used, similarly for cis analysis. As expected, we find that the contact conservation is higher for Human-Mouse (AUC > 0.8) when compared with Human-Fly (AUC 0.5) or Mouse-Fly (AUC 0.5) (Figure 3C) in both cis and trans. We also find that genes with high contact conservation in Human-Mouse are likely to have high contact coexpression.

We also re-validated the power of the meta-3C network: we compared the ‘contact conservation’ scores for individual and meta-3C networks at various resolutions. We reassuringly find that the meta-3C network outperforms individual projects at high resolutions in both cis and trans (Figure 3D, S4C and D). This again suggests that the meta-3C network is efficient in capturing chromatin contacts when compared to individual networks. Although the conservation of cis chromatin structure across species is not surprising and is evident in the presence of syntenic regions between species, the conservation of trans-chromatin contacts is noteworthy. It suggests that the trans-chromatin structure is likely selected for preservation to maintain function.

We further investigated the evolution of trans chromatin contacts in Human by comparing the degree to which the Human contacts can predict coexpression across several species. This method allowed us to extend our analysis to species for which the meta-3C network is not available. Each gene’s neighborhood is defined by ranking all edges in the chromatin contact network of one species and then used to predict the gene’s top 10% of coexpressed gene pairs in another species. We call this “contact coexpression conservation” and calculate the AUC as above. When contact coexpression conservation is plotted along with the phylogenetic distance across species, we find that the performance decreases with an increase in phylogenetic distance using both cis and trans meta-3C networks (Figure 3F). This suggests that the contacts diverge as the species pair becomes distant across evolution. The number of 1:1 orthologs also decreases with an increase in the phylogenetic distance (Figure S4B) and it seemed possible that our observation was dominated by the number of ortholog pairs between species. To eliminate this possibility, we redid our analysis but using only the same set of ortholog genes (429 genes) in each species and our result persisted (Figure 3G). Species more than 100 million years of distance (mya) from Humans have stronger divergence in contacts when compared to species within the Mammalia Class. The species included in Figure 3G were limited to the Chordata phylum to ensure a reasonable number of genes in the analysis.

### Data availability and Online Tool

In order to facilitate the broad adoption of meta-3C by the community, we have made data available via an online tool (https://gillisweb.cshl.edu/HiC/). Contact data can be obtained in two ways: a] Network download: Direct download of the desired resolution, species meta-3C contact matrix in cis or trans available at https://labshare.cshl.edu/shares/gillislab/resource/HiC in HiCMatrix format (https://github.com/deeptools/HiCMatrix). b] Gene vector download: Contact frequency with every genomic loci at the chosen resolution and for any desired gene found in the respective species (Figure 4A). The downloaded file is in bed file format which can be uploaded to UCSC genome browser for further analysis as desired (Figure 4B).

**Fig. 4.**
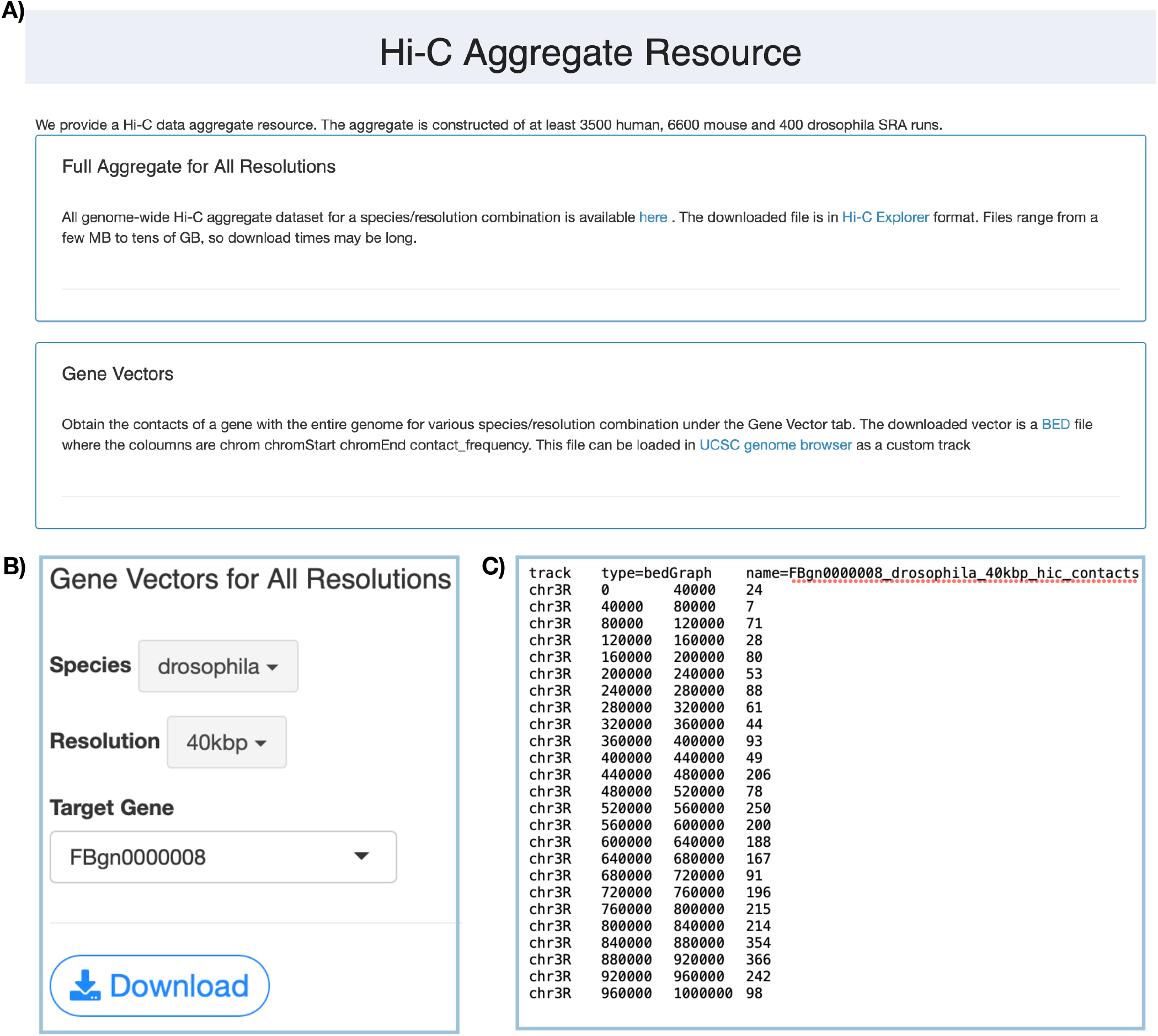
Snapshot of the tool page (https://gillisweb.cshl.edu/HiC/) (A) gene vector download page (B). C) Downloaded bed file snapshot where the first column is chromosome name, next two columns are genomic bin and the last column is raw contact frequency.

## Discussion

In this work, we created a high-depth, genome-wide chromatin contact map using a meta-analytic approach, validated it, and further used it to reveal chromatin structure to function relationships. We find that for the three species analyzed in this study (Human, Mouse, and Fly), chromatin contacts strongly predicted the coexpression of genes. We also show that chromatin contacts are better than linear proximity for predicting eQTLs when high-resolution chromatin contact data is available. Our results persist even when only long-range chromatin contacts are analyzed. Additionally, we find that trans chromosomal contacts show evidence of conservation across species.

Meta-3C networks are an effective means for capturing otherwise hard to characterize long-range interactions providing potentially uniquely important practical applications. One important application for a wide area of genomics is their ability to prioritize distant target genes for variants. We expect these networks to be powerful training data for future machine learning attempts to predict chromosomal contacts, an important area of ongoing research (10–13). Additionally, meta-3C networks can be used with other cell-type-specific ‘omics datasets such as ChIP-Seq to reveal cell-type-specific enhancer-promoter contacts. Previously, Nasser et al (26, 27) used averaged Hi-C data across 10 cell-types in their ABC model to accurately make cell-type-specific enhancer–gene predictions. Thus, the continuing evolution of methods with improved specificity is likely to complement our better-powered but less condition-specific meta-analytic approach.

Within 3C analysis, and even outside of it, aggregation of data is well appreciated to be a useful strategy. Reproducible biological replicates within the same study are often combined to increase the density of 3C data thereby capturing more interactions (15, 28). Our approach can be thought of as the most extreme version of this idea, combining experiments as broadly as possible to capture statistical relationships that are common. This is most useful if the depth is a major limitation, as in trans contacts, as it comes with the cost of a loss of condition-specificity. Thus, the route forward for the field as a whole will doubtless involve both improved specificity, integration, and interpretive methods.

In summary, our study sheds new light on the functional role of long-range and trans chromosomal contacts and provides a critical resource for use by a wide range of genomics research.

## Materials and Methods

### 3C Data Sources and processing pipeline

The 3C data for each species were obtained from SRA search (https://www.ncbi.nlm.nih.gov/sra/) with the field limitations of “Organism”: [“Homo sapiens”, “Mus musculus”, “Drosophila melanogaster”], “Strategy”: “hi c”. We found 3913, 8431, 502 samples for Human, Mouse, and Fly respectively. We also added 268, 17, and 25 samples manually that were labeled OTHER in SRA, but were deemed to be valid 3C data based on publication details. After manual additions, filtering out Runs without available restriction enzyme information and excluding Runs that failed processing, we had 3621, 6733, 487 samples for Human, Mouse, and Fly. In total, we aggregated 119 Human Projects, 33 Mouse Projects, and 29 Fly Projects (Table S1, S2, and S3). All samples were reprocessed from short read sequence data to reduce differential computational noise across experiments. Restriction enzymes were identified for each sample from the literature.

SRA files were downloaded using prefetch, then converted to paired FASTQ files using fasterq-dump. FASTQ files were processed using the HiCUP tool (29), with the alteration that short reads were aligned using the STAR aligner, instead of the default Bowtie2. HiCUP truncates the reads based on restriction site, aligns them, and filters artifactual and duplicated data. Reads were aligned to the hg38, mm10, and dm6 genomes. Output SAM files were converted to indexed and compressed Pairs files using the bam2pairs tool. Finally, pairwise chromosome-chromosome contact matrices were generated at single base-pair resolution.

### Building CC-maps

To obtain CC-maps, each chromosome is divided up into “bins” of a specific size. The number of base pairs in each bin represents the “resolution” of the matrix. The contact frequency for each bin pair is obtained by summing the reads falling in that bin. CC-maps were generated at 8 resolutions (1KB, 5KB, 10KB, 25KB, 40KB, 100KB, 250KB, and 500KB) in cis for all species and trans for only Fly. For Human and Mouse, trans CC-map at 1KB and 5KB resolutions were not processed due to high memory requirements (more than 2TB). These files were written in HiCMatrix (https://github.com/deeptools/HiCMatrix) h5 format. For each species, we excluded sex chromosomes and considered only autosomes (Human: chr1 to chr22, Mouse:chr1 to chr19 and Fly: chr2L, chr2R, chr3L, chr3R, chr4). The contact frequency of each genomic pair coordinate was summed across runs to generate project-level CC-maps. The sequencing depth of a project in cis and trans is obtained by summing all the contacts in cis and trans respectively. The contact frequency was KR-normalized separately for the cis and trans networks to adjust for nonuniformities in coverage introduced due to experimental bias (30) using hicCorrectMatrix tool of HiCexplorerV3.6 (31). All project level CC-maps within each species were further summated to create species level meta-3C maps. To determine contact frequency between each gene pair we use the maximum contact frequency between each bin in which genes reside. Gene TSS and TES were used to determine the bins in which the gene resides. List of genes, TSS, and TES were obtained as GTF files from ENSEMBEL (September 2019). A list of 1:1 orthologs for pair of species was obtained from OrthoDB (32). Species diverge time was sourced from Timetree (33). Gene compartments were identified using ‘hicPCA’ tool of HiCexplorerV3.6 (31) using each chromosome KR-normalized cis-contact matrix.

### Coexpression data

The coexpression network used in this study is a ‘high confidence gene’ aggregated coexpression network generated using the method previously described in CoCoCoNet (20). In brief, several bulk RNA-seq datasets were obtained from NCBI’s SRA database (unique SRA Study IDs). Networks for each dataset are built by calculating the Spearman correlation between all pairs of genes, then ranking the correlation coefficients for all gene-gene pairs, with NAs assigned the median rank. Each network is then rank standardized and normalized by dividing through by the maximum rank. Aggregate networks are then generated by averaging rank standardized networks from individual datasets.

### eQTL data source and processing

A list of tissue-specific ‘significant’ variant gene pair associations and ‘all’ variant gene pair associations (including non-significant associations) across 54 tissues along with the distance between the variant and gene TSS (at bp resolution) were obtained from GTEx Portal v8 at https://gtexportal.org. Since the meta-3C network is not tissue-specific, we combined the data across tissues to generate a set of unique ‘significant’ and ‘all’ variant gene pair associations. To obtain a list of ‘non-significant’ gene pair associations, ‘significant’ variant gene pair associations were removed from ‘all’ variant gene pair associations data. All variants in the coding regions and up to 1KB of any gene TSS and TES were removed. For performance score 1KB cis CC-map is used and for each eVariant only genes in unique bins are tested.

## Data Availability

The Meta 3C networks for Human, Mouse, and Fly are available for download from online tool (https://gillisweb.cshl.edu/HiC/) or direct download (https://labshare.cshl.edu/shares/gillislab/resource/HiC/)

## Supplementary Information

**Figures**

Fig S1: B) C) D) and E) of Fig 2 for Mouse

Fig S2: B) C) D) and E) of Fig 2 for Fly

Fig S3: Compartment vs contact prediction in trans

Fig S4: Fig 3 B) in cis and cross-species conservation at various resolutions

**Tables**

Table S1: Details of each individual project used for building Human meta-3C network

Table S2: Same as Table S1 but for Mouse:

Table S3: Same as Table S1 but for Fly

**Table S1.** Details of each individual project used for building Human meta-3C network.

| Project | Reference study | of runs | Total Contacts |
| --- | --- | --- | --- |
| SRP050102 | (15) | 217 | 21903231337 |
| SRP012412 | (34) | 618 | 21526850534 |
| SRP152979 | (35) | 89 | 22953596600 |
| SRP152879 | (36) | 27 | 7171511913 |
| SRP118999 | (37) | 501 | 12590883331 |
| SRP154953 | (38) | 31 | 9373339255 |
| SRP199098 | (39) | 8 | 10496943998 |
| SRP234115 | (40) | 29 | 8756864842 |
| SRP094854 |  | 39 | 11764162990 |
| SRP149906 | (36) | 41 | 5433846272 |
| SRP165933 | (41) | 20 | 8622402152 |
| SRP212226 | (11) | 59 | 5567660318 |
| SRP117084 | (42) | 136 | 9677237142 |
| SRP218691 |  | 47 | 11527982949 |
| SRP233368 | (43) | 16 | 6174138589 |
| SRP106040 | (44) | 14 | 3806131390 |
| SRP125488 |  | 9 | 3406425709 |
| SRP250432 |  | 8 | 3818798951 |
| SRP141473 | (45) | 37 | 3012977379 |
| ERP107279 |  | 12 | 4052904958 |
| SRP224133 | (46) | 23 | 8288729106 |
| SRP184300 |  | 8 | 3236851964 |
| SRP168606 | (24) | 15 | 5024303634 |
| SRP216194 | (47) | 63 | 4764862118 |
| SRP178527 | (48) | 3 | 3173762302 |
| SRP173234 | (49) | 18 | 3047832982 |
| SRP239849 | (50) | 17 | 3838499161 |
| SRP106379 |  | 18 | 4505448986 |
| SRP162098 | (51) | 14 | 2758854779 |
| SRP120957 |  | 14 | 1880560283 |
| DRP005280 |  | 3 | 1138767416 |
| SRP150259 | (52) | 9 | 1038770389 |
| SRP170743 |  | 10 | 2670689487 |
| SRP131003 | (53) | 14 | 2249959398 |
| SRP158113 |  | 7 | 2746843677 |
| SRP186190 |  | 8 | 1364868796 |
| SRP212073 | (54) | 10 | 1901334229 |
| SRP133031 | (55) | 30 | 676189370 |
| SRP135798 |  | 10 | 1791532495 |
| SRP131871 | (56) | 54 | 2247326065 |
| SRP158276 | (57) | 9 | 1345162772 |
| SRP114754 |  | 20 | 1199618716 |
| SRP267107 | (58) | 36 | 3093708888 |
| SRP227918 | (59) | 4 | 2007602381 |
| SRP271101 | (60) | 5 | 1475289101 |
| SRP186277 |  | 8 | 653828297 |
| SRP115913 | (61) | 25 | 4731907294 |
| SRP157799 | (62) | 16 | 1458459569 |
| SRP110964 |  | 9 | 2237670786 |
| SRP194362 | (63) | 10 | 1553016593 |
| SRP151075 | (64) | 3 | 1400791640 |
| SRP157894 | (65) | 4 | 1878237731 |
| SRP160101 |  | 17 | 1215405010 |
| SRP157048 | (66) | 6 | 1781845628 |
| SRP221518 | (67) | 38 | 2296723083 |
| SRP225696 | (68) | 8 | 683636873 |
| ERP104251 |  | 4 | 774712749 |
| SRP105082 | (69) | 699 | 891951555 |
| SRP223060 | (70) | 9 | 1575825293 |
| SRP234897 | (71) | 32 | 1072337563 |
| SRP250333 | (72) | 16 | 2535101216 |
| SRP113633 | (73) | 6 | 1123994483 |
| SRP186012 | (74) | 12 | 865590451 |
| SRP199225 | (75) | 16 | 568841794 |
| SRP107176 | (76) | 2 | 825826168 |
| SRP105181 | (77) | 3 | 384767159 |
| SRP066852 |  | 4 | 394762758 |
| SRP095110 |  | 2 | 303999589 |
| SRP162056 |  | 5 | 670529013 |
| SRP201909 | (78) | 8 | 694131646 |
| SRP153415 | (79) | 4 | 574542701 |
| SRP127183 | (80) | 2 | 624271125 |
| ERP118600 |  | 3 | 804408269 |
| SRP274139 | (81) | 3 | 564434221 |
| SRP115572 | (82) | 76 | 372409309 |
| SRP099610 | (83) | 4 | 620448038 |
| SRP108500 | (84) | 60 | 411838736 |
| SRP195614 | (85) | 4 | 371304283 |
| SRP235557 | (86) | 1 | 607333953 |
| SRP264796 | (87) | 3 | 630348911 |
| SRP197114 |  | 6 | 266221302 |
| SRP132233 | (88) | 1 | 456935896 |
| SRP244334 |  | 2 | 360094743 |
| SRP113478 |  | 5 | 131412179 |
| SRP107148 |  | 2 | 243243433 |
| SRP083971 |  | 2 | 216849003 |
| SRP192392 | (89) | 2 | 417818971 |
| SRP145420 | (90) | 6 | 359393402 |
| SRP152361 | (91) | 2 | 364093153 |
| SRP141229 |  | 5 | 359844610 |
| SRP261300 |  | 2 | 140068804 |
| SRP154986 | (92) | 3 | 262627353 |
| SRP261299 |  | 2 | 190528229 |
| SRP150629 | (93) | 10 | 201563278 |
| SRP111140 |  | 1 | 205355109 |
| SRP130935 | (94) | 8 | 511638458 |
| SRP165232 |  | 2 | 176738015 |
| SRP098826 |  | 2 | 146110482 |
| SRP102403 | (95) | 2 | 313084130 |
| SRP154399 |  | 2 | 184474817 |
| SRP090318 | (96) | 4 | 163096218 |
| DRP005173 | (97) | 9 | 237678253 |
| SRP107149 |  | 2 | 168664932 |
| SRP245657 |  | 2 | 213200567 |
| SRP149124 |  | 4 | 369028739 |
| SRP060755 |  | 2 | 136528125 |
| SRP163366 | (98) | 1 | 35514017 |
| SRP071243 |  | 1 | 151366598 |
| SRP272124 | (99) | 1 | 101271490 |
| SRP103077 |  | 2 | 128669702 |
| SRP163908 |  | 2 | 149842370 |
| SRP170855 | (100) | 1 | 136131087 |
| SRP217227 | (74) | 2 | 63146633 |
| SRP132876 | (101) | 1 | 74105800 |
| SRP100408 | (102) | 10 | 32147936 |
| SRP105086 |  | 1 | 17034573 |
| SRP076397 |  | 2 | 990958 |
| SRP182670 |  | 1 | 352599 |
| SRP109036 |  | 4 | 9968 |

**S1.**
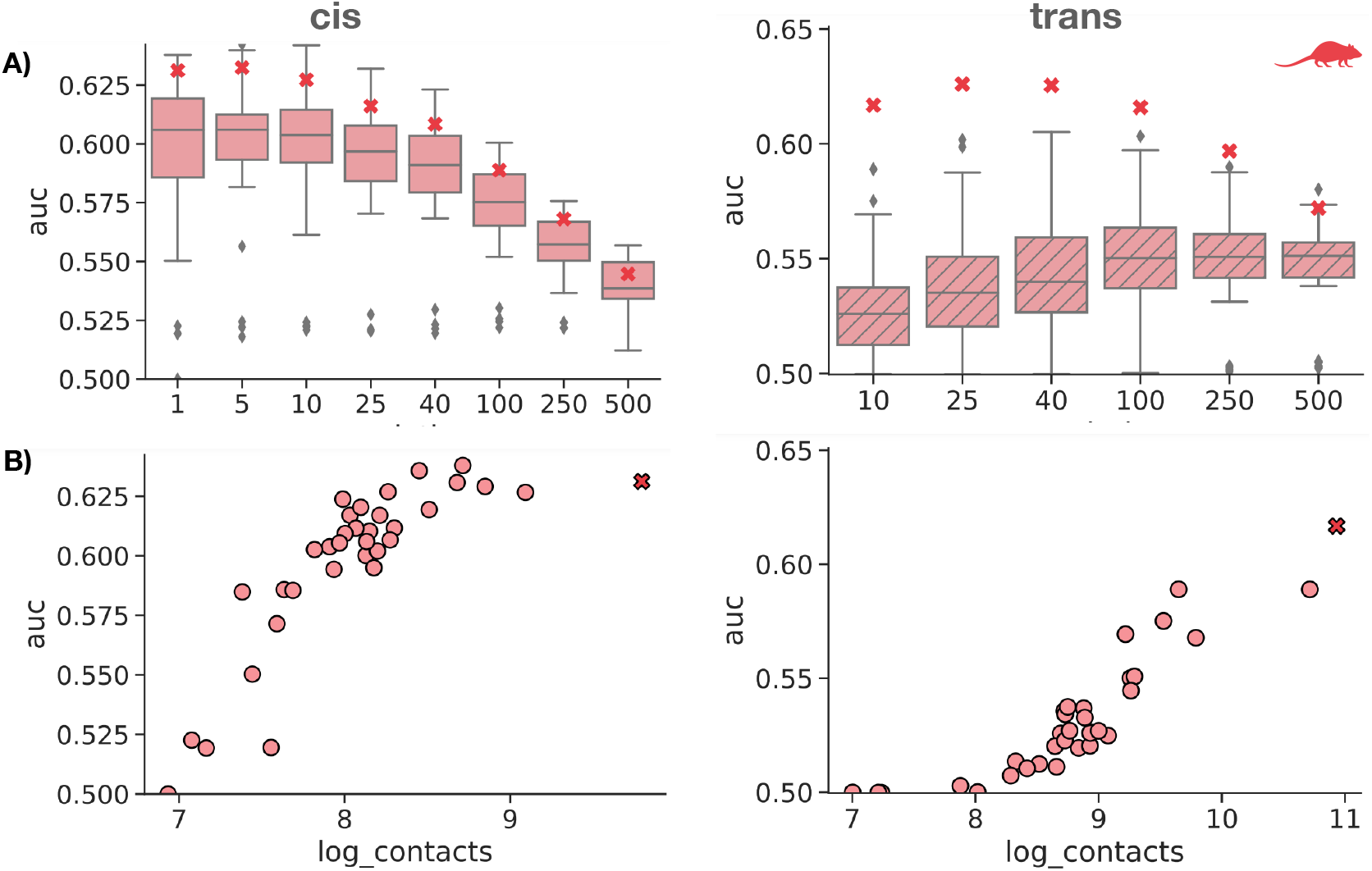
A) The boxplot shows the distribution of median AUC across all genes for each project at various resolutions in cis (left) and trans (right). B) The circles represent the median AUC across all genes for each project vs sequencing depth at 1KB resolution in cis (left) and 10KB resolution in trans (right).

**S2.**
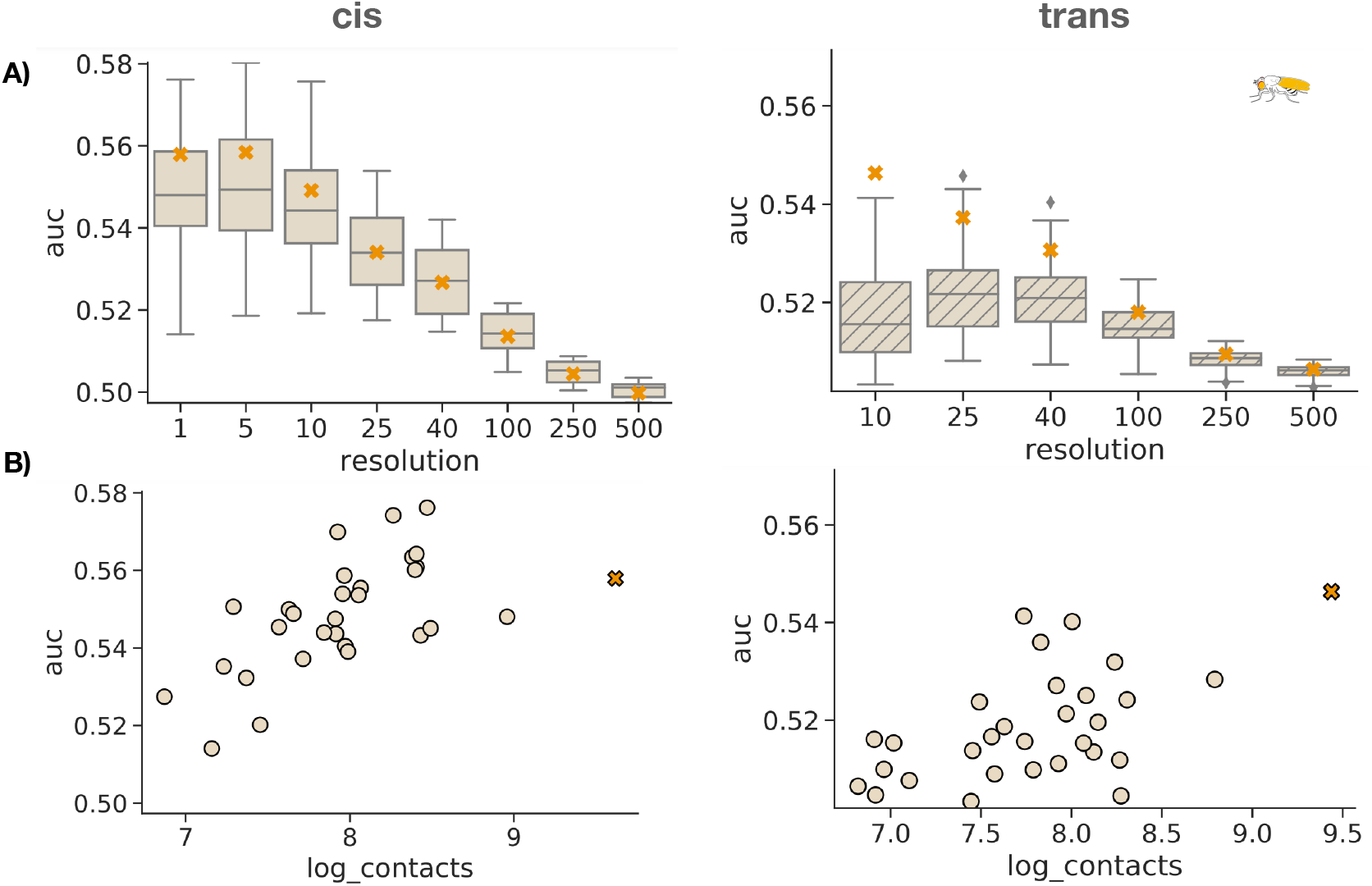
A) The boxplot shows the distribution of median AUC across all genes for each project at various resolutions in cis (left) and trans (right). B) The circles represent the median AUC across all genes for each project vs sequencing depth at 1KB resolution in cis (left) and 10KB resolution in trans (right).

**S3.**
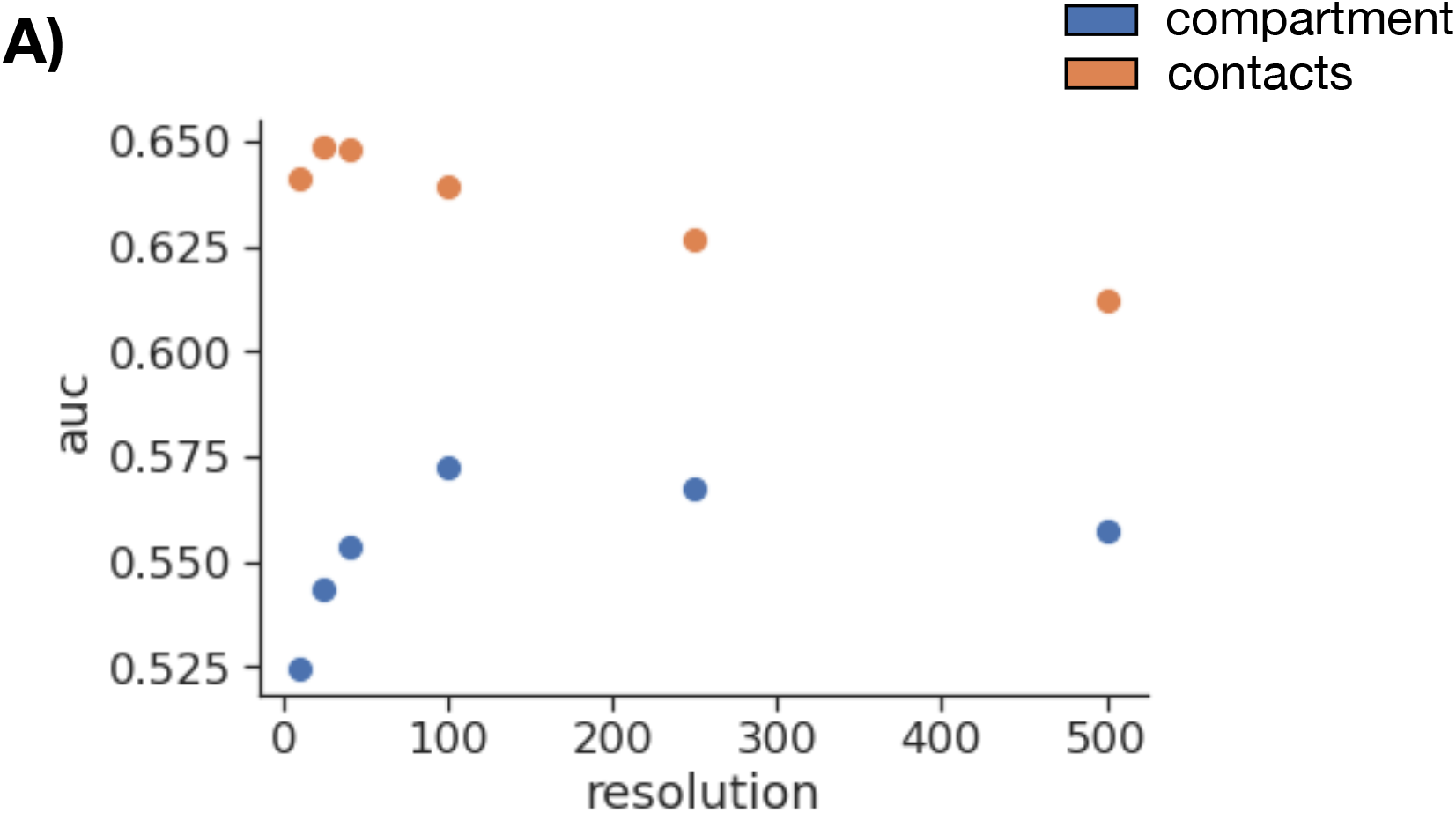
Performance of contact co-expression compartment co-expression at various resolutions.

**S4.**
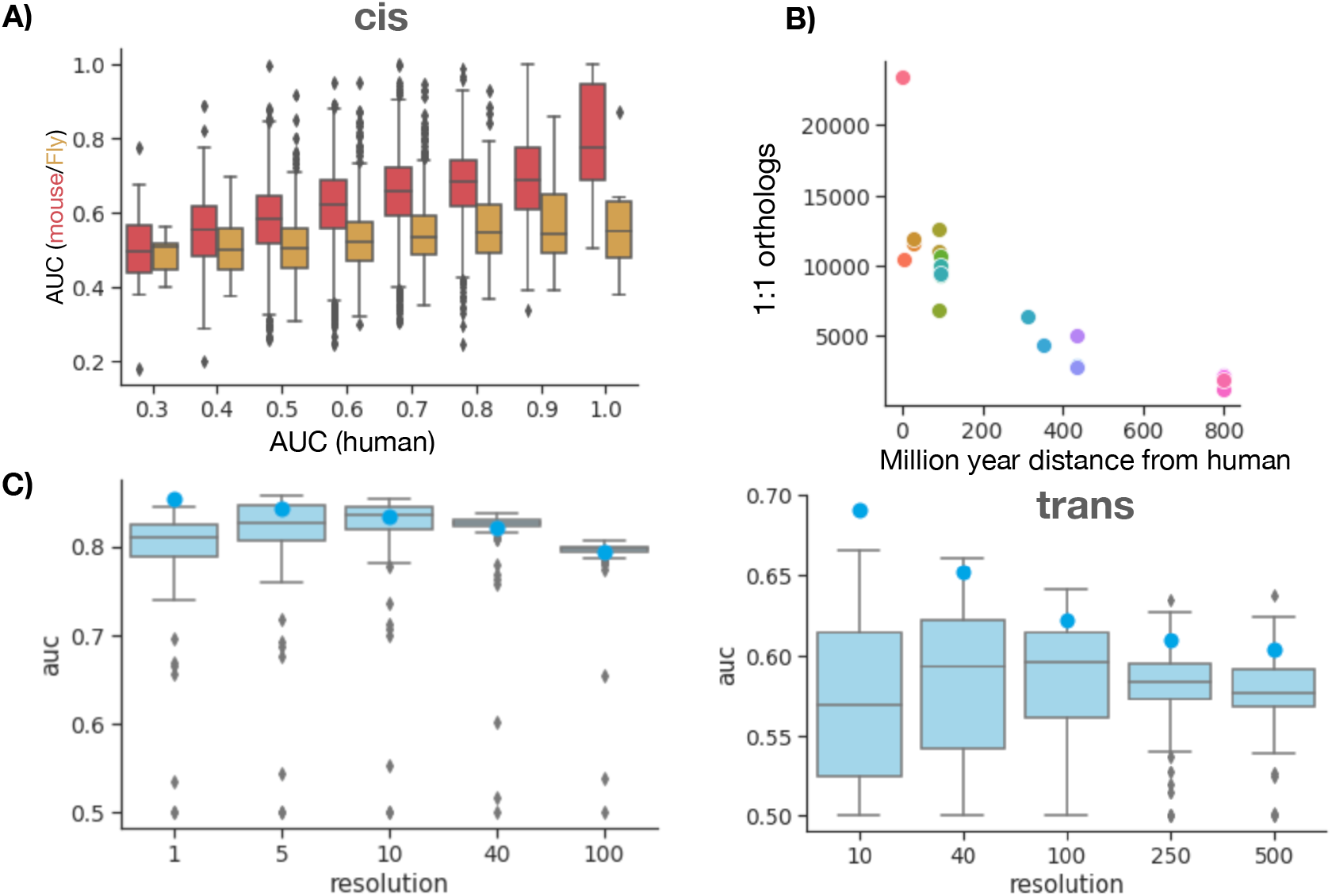
Chromatin contacts are conserved across species. A) Performance of gene contact to predict coexpression in Human vs other species (Mouse and Fly) in cis. B) Number of 1:1 orthologs between human and various other species. C) The boxplot shows the distribution of median AUC across all genes for each project at various resolutions in cis (left) and trans (right).

**Table S2.** Details of each individual project used for building Mouse meta-3C network.

| Project | Reference study | of runs | Total Contacts |
| --- | --- | --- | --- |
| SRP217487 | (103) | 573 | 72472272945 |
| SRP101928 | (104) | 56 | 19175842719 |
| SRP075985 | (105) | 573 | 13165889801 |
| SRP105082 | (69) | 361 | 11550001954 |
| SRP165933 | (41) | 22 | 8027513684 |
| SRP261290 | (106) | 20 | 5039895689 |
| SRP118601 | (107) | 200 | 3677569528 |
| SRP107774 | (108, 109) | 60 | 6606308156 |
| SRP226118 |  | 68 | 4301950689 |
| SRP252213 | (110) | 24 | 2762736772 |
| SRP229756 |  | 52 | 3844845689 |
| SRP250878 |  | 56 | 3338952796 |
| SRP131117 | (111) | 21 | 3373187973 |
| SRP247488 | (112) | 52 | 2550365588 |
| SRP223513 | (113) | 78 | 4143881490 |
| SRP119332 | (114) | 37 | 2279482102 |
| SRP268173 | (67) | 36 | 2920915986 |
| SRP270993 | (115) | 57 | 2276128337 |
| SRP179647 | (116) | 48 | 2443326437 |
| SRP255620 | (117) | 22 | 1699512024 |
| SRP100871 | (118) | 30 | 3309838005 |
| SRP192917 | (11) | 35 | 3455012039 |
| SRP156597 | (119) | 63 | 3274402425 |
| SRP227097 | (67) | 37 | 1925624481 |
| ERP114475 |  | 24 | 1092282117 |
| SRP249897 | (120) | 2544 | 1171971517 |
| SRP096571 | (121) | 48 | 641165329 |
| SRP144391 | (122) | 52 | 930492853 |
| SRP110616 | (123) | 211 | 790590876 |
| SRP292639 | (120) | 1002 | 598654244 |
| SRP194410 | (124) | 67 | 252418461 |
| SRP200567 | (125) | 131 | 307721826 |
| SRP218950 | (119) | 73 | 173739328 |

**Table S3.** Details of each individual project used for building Mouse meta-3C network.

| Project | Reference study | of runs | Total Contacts |
| --- | --- | --- | --- |
| ERP122732 | (126) | 76 | 4268124240 |
| SRP165773 | (127) | 12 | 1214638382 |
| SRP119928 | (128) | 15 | 485650828 |
| ERP112882 |  | 38 | 509996238 |
| SRP050096 | (129) | 17 | 1176311012 |
| SRP223221 | (130) | 4 | 473122715 |
| SRP097891 | (131) | 13 | 500368469 |
| ERP016479 |  | 17 | 1215808351 |
| SRP104256 | (132) | 11 | 565652294 |
| SRP186730 | (133) | 8 | 1284274100 |
| SRP230396 | (134) | 32 | 1330184944 |
| SRP107636 | (135) | 10 | 255856377 |
| SRP073988 | (136) | 6 | 1035207740 |
| SRP111713 | (137) | 6 | 1030632363 |
| SRP107637 | (135) | 7 | 207440463 |
| SRP195621 | (85) | 4 | 493068117 |
| SRP193880 |  | 24 | 780569203 |
| SRP168946 | (138) | 4 | 406168278 |
| SRP107556 | (135) | 2 | 414118766 |
| SRP158369 | (139) | 8 | 368378383 |
| SRP110166 |  | 10 | 424475815 |
| SRP165772 | (140) | 6 | 232779025 |
| ERP112723 |  | 12 | 80844400 |
| SRP219433 |  | 6 | 81129339 |
| SRP156199 | (141) | 8 | 184854440 |
| SRP199618 | (142) | 122 | 96407920 |
| SRP132075 | (143) | 4 | 119651337 |
| SRP140881 |  | 4 | 85005475 |
| SRP181908 | (144) | 1 | 36788399 |

## Notes

https://gillisweb.cshl.edu/HiC/

### Competing Interest Statement

The authors have declared no competing interest.

### Summary of Updates

Added Fig S3 to comment on the role of contacts vs chromatin compartment in predicting coexpression. Added tables S1, S2 and S3

## References

1. Diament A, Tuller T (2019) Modeling three-dimensional genomic organization in evolution and pathogenesis. Seminars in Cell Developmental Biology 90:78–93.

2. Delaneau O, et al. (2019) Chromatin three-dimensional interactions mediate genetic effects on gene expression. Science 364(6439).

3. Xu H, Zhang S, Yi X, Plewczynski D, Li MJ (2020) Exploring 3D chromatin contacts in gene regulation: The evolution of approaches for the identification of functional enhancer-promoter interaction.

4. Sarnataro S, Chiariello AM, Esposito A, Prisco A, Nicodemi M (2017) Structure of the human chromosome interaction network. PLoS One 12(11):e0188201.

5. Dekker J, Misteli T (2015) Long-Range chromatin interactions. Cold Spring Harb. Perspect. Biol. 7(10):a019356.

6. Maass PG, Barutcu AR, Rinn JL (2019) Interchromosomal interactions: A genomic love story of kissing chromosomes. J. Cell Biol. 218(1):27–38.

7. Lieberman-Aiden E, et al. (2009) Comprehensive mapping of long-range interactions reveals folding principles of the human genome. Science 326(5950):289–293.

8. Fang R, et al. (2016) Mapping of long-range chromatin interactions by proximity ligation-assisted ChIP-seq. Cell Res. 26(12):1345–1348.

9. Dryden NH, et al. (2014) Unbiased analysis of potential targets of breast cancer susceptibility loci by capture Hi-C. Genome Res. 24(11):1854–1868.

10. Fudenberg G, Kelley DR, Pollard KS (2020) Predicting 3D genome folding from DNA sequence with akita. Nat. Methods 17(11):1111–1117.

11. Zhang J, et al. (2019) Spatial clustering and common regulatory elements correlate with coordinated gene expression. PLoS Computational Biology 15(3):e1006786.

12. Zhang Y, et al. (2018) Enhancing Hi-C data resolution with deep convolutional neural network HiCPlus. Nature Communications 2018 9:1 9(1):1–9.

13. Schwessinger R, et al. (2020) DeepC: predicting 3D genome folding using megabase-scale transfer learning. Nat. Methods 17(11):1118–1124.

14. Ay F, Bailey TL, Noble WS (2014) Statistical confidence estimation for Hi-C data reveals regulatory chromatin contacts. Genome Res. 24(6):999–1011.

15. Rao SS, et al. (2014) A 3D map of the human genome at kilobase resolution reveals principles of chromatin looping. Cell 159(7):1665–1680.

16. Bulathsinghalage C, Liu L (2020) Network-based method for regions with statistically frequent interchromosomal interactions at single-cell resolution. BMC Bioinformatics 21(Suppl 14):369.

17. Xiong K, Ma J (2019) Revealing Hi-C subcompartments by imputing inter-chromosomal chromatin interactions. Nat. Commun. 10(1):5069.

18. Varrone M, Nanni L, Ciriello G, Ceri S (2020) Exploring chromatin conformation and gene co-expression through graph embedding. Bioinformatics 36(Supplement_2):i700–i708.

19. Babaei S, et al. (2015) Hi-C Chromatin Interaction Networks Predict Co-expression in the Mouse Cortex. PLoS Comput. Biol. 11(5):e1004221.

20. Lee J, Shah M, Ballouz S, Crow M, Gillis J (2020) CoCoCoNet: conserved and comparative co-expression across a diverse set of species. Nucleic Acids Research 48(W1):W566–W571.

21. Wang H, Yang J, Zhang Y, Wang J (2021) Discover novel disease-associated genes based on regulatory networks of long-range chromatin interactions. Methods 189:22–33.

22. Kong N, Jung I (2020) Long-range chromatin interactions in pathogenic gene expression control. Transcription 11(5):211–216.

23. Stacey D, et al. (2019) ProGeM: a framework for the prioritization of candidate causal genes at molecular quantitative trait loci. Nucleic Acids Res. 47(1):e3.

24. Eres IE, Luo K, Hsiao CJ, Blake LE, Gilad Y (2019) Reorganization of 3D genome structure may contribute to gene regulatory evolution in primates. PLoS Genetics 15(7):e1008278.

25. Krefting J, Andrade-Navarro MA, Ibn-Salem J (2018) Evolutionary stability of topologically associating domains is associated with conserved gene regulation. BMC Biology 2018 16:1 16(1):1–12.

26. Nasser J, et al. (2021) Genome-wide enhancer maps link risk variants to disease genes. Nature 593(7858):238–243.

27. Fulco CP, et al. (2019) Activity-by-contact model of enhancer-promoter regulation from thousands of CRISPR perturbations. Nat. Genet. 51(12):1664–1669.

28. Won H, et al. (2016) Chromosome conformation elucidates regulatory relationships in developing human brain. Nature 538(7626):523–527.

29. Wingett S, et al. (2015) HiCUP: pipeline for mapping and processing Hi-C data. F1000Res. 4:1310.

30. Knight PA, Ruiz D (2012) A fast algorithm for matrix balancing. IMA J. Numer. Anal. 33(3):1029–1047.

31. Wolff J, et al. (2020) Galaxy HiCExplorer 3: a web server for reproducible Hi-C, capture Hi-C and single-cell Hi-C data analysis, quality control and visualization. Nucleic Acids Res. 48(W1):W177–W184.

32. Kriventseva EV, et al. (2019) OrthoDB v10: sampling the diversity of animal, plant, fungal, protist, bacterial and viral genomes for evolutionary and functional annotations of orthologs. Nucleic Acids Res. 47(D1):D807–D811.

33. Kumar S, Stecher G, Suleski M, Hedges SB (2017) TimeTree: A resource for timelines, timetrees, and divergence times. Mol. Biol. Evol. 34(7):1812–1819.

34. Heidari N, et al. (2014) Genome-wide map of regulatory interactions in the human genome. Genome Research 24(12):1905.

35. Arvanitis M, et al. (2020) Genome-wide association and multi-omic analyses reveal ACTN2 as a gene linked to heart failure. Nat. Commun. 11(1):1122.

36. Lu L, et al. (2020) Robust Hi-C maps of Enhancer-Promoter interactions reveal the function of non-coding genome in neural development and diseases. Mol. Cell 79(3):521–534.e15.

37. Rao SSP, et al. (2017) Cohesin loss eliminates all loop domains.

38. Lawlor N, et al. (2019) Multiomic profiling identifies cis-regulatory networks underlying human pancreatic *β* cell identity and function. Cell Rep. 26(3):788–801.e6.

39. Ochi Y, et al. (2020) Combined Cohesin-RUNX1 deficiency synergistically perturbs chromatin looping and causes myelodysplastic syndromes. Cancer Discov. 10(6):836–853.

40. Stik G, et al. (2020) CTCF is dispensable for immune cell transdifferentiation but facilitates an acute inflammatory response. Nat. Genet. 52(7):655–661.

41. El Khattabi L, et al. (2019) A pliable mediator acts as a functional rather than an architectural bridge between promoters and enhancers. Cell 178(5):1145–1158.e20.

42. Heinz S, et al. (2018) Transcription elongation can affect genome 3D structure. Cell 174(6):1522–1536.e22.

43. Kang H, et al. (2020) Dynamic regulation of histone modifications and long-range chromosomal interactions during postmitotic transcriptional reactivation. Genes Dev. 34(13-14):913–930.

44. Zirkel A, et al. (2017) Topological demarcation by HMGB2 is disrupted early upon senescence entry across cell types and induces CTCF clustering.

45. Song M, et al. (2019) Mapping cis-regulatory chromatin contacts in neural cells links neuropsychiatric disorder risk variants to target genes. Nat. Genet. 51(8):1252–1262.

46. Wutz G, et al. (2020) ESCO1 and CTCF enable formation of long chromatin loops by protecting cohesinSTAG1 from WAPL.

47. Kloetgen A, et al. (2020) Three-dimensional chromatin landscapes in T cell acute lymphoblastic leukemia. Nat. Genet. 52(4):388–400.

48. Liu L, Li QZ, Jin W, Lv H, Lin H (2019) Revealing Gene Function and Transcription Relationship by Reconstructing Gene-Level Chromatin Interaction. Computational and Structural Biotechnology Journal 17:195–205.

49. Nir G, et al. (2018) Walking along chromosomes with super-resolution imaging, contact maps, and integrative modeling. PLoS Genet. 14(12):e1007872.

50. Yang J, et al. (2020) Analysis of chromatin organization and gene expression in T cells identifies functional genes for rheumatoid arthritis. Nat. Commun. 11(1):4402.

51. Ferrari R, et al. (2020) TFIIIC binding to alu elements controls gene expression via chromatin looping and histone acetylation. Mol. Cell 77(3):475–487.e11.

52. Jacobson EC, et al. (2018) Migration through a small pore disrupts inactive chromatin organization in neutrophil-like cells.

53. Dily FL, et al. (year?) Hormone control regions mediate opposing steroid receptor-dependent genome organizations.

54. Zhang X, et al. (2021) The loss of heterochromatin is associated with multiscale three-dimensional genome reorganization and aberrant transcription during cellular senescence. Genome Res.

55. Rodriguez A (2021) High HDL-Cholesterol paradox: SCARB1-LAG3-HDL axis. Curr. Atheroscler. Rep. 23(1):5.

56. Paulsen J, et al. (2019) Long-range interactions between topologically associating domains shape the four-dimensional genome during differentiation.

57. Achinger-Kawecka J, et al. (2020) Epigenetic reprogramming at estrogen-receptor binding sites alters 3D chromatin landscape in endocrine-resistant breast cancer. Nat. Commun. 11(1):320.

58. Mitter M, et al. (2020) Conformation of sister chromatids in the replicated human genome. Nature 586(7827):139–144.

59. Senigl F, et al. (2019) Topologically associated domains delineate susceptibility to somatic hypermutation. Cell Rep. 29(12):3902–3915.e8.

60. Guo D, et al. (2021) Synergistic alterations in the multilevel chromatin structure anchor dysregulated genes in small cell lung cancer. Comput. Struct. Biotechnol. J. 19:5946–5959.

61. Wutz G, et al. (2017) Topologically associating domains and chromatin loops depend on cohesin and are regulated by CTCF, WAPL, and PDS5 proteins. EMBO J. 36(24):3573–3599.

62. Iwasaki O, et al. (2019) Involvement of condensin in cellular senescence through gene regulation and compartmental reorganization. Nat. Commun. 10(1):5688.

63. Dileep V, et al. (2019) Rapid irreversible transcriptional reprogramming in human stem cells accompanied by discordance between replication timing and chromatin compartment. Stem Cell Reports 13(1):193–206.

64. Tian L, et al. (2018) Abstract 1485: Allelic specificity of immunoglobulin heavy chain (IGH) translocation in b-cell acute lymphoblastic leukemia (B-ALL) unveiled by long-read sequencing. Cancer Res. 78(13_Supplement):1485–1485.

65. Guo Y, et al. (2018) CRISPR-mediated deletion of prostate cancer risk-associated CTCF loop anchors identifies repressive chromatin loops. Genome Biol. 19(1):160.

66. Ooi WF, et al. (2020) Integrated paired-end enhancer profiling and whole-genome sequencing reveals recurrent CCNE1 and IGF2 enhancer hijacking in primary gastric adenocarcinoma. Gut 69(6):1039–1052.

67. Zhang Y, Cao J (2020) GSimPy: A Python package for measuring group similarity. SoftwareX 12:100526.

68. Hyle J, et al. (2019) Acute depletion of CTCF directly affects MYC regulation through loss of enhancer–promoter looping. Nucleic Acids Res. 47(13):6699–6713.

69. Vian L, et al. (2018) The energetics and physiological impact of cohesin extrusion. Cell 173(5):1165.

70. Ray-Jones H, et al. (2020) Mapping DNA interaction landscapes in psoriasis susceptibility loci highlights KLF4 as a target gene in 9q31. BMC Biol. 18(1):47.

71. Choudhary MN, et al. (2020) Co-opted transposons help perpetuate conserved higher-order chromosomal structures. Genome Biol. 21(1):16.

72. Thiecke MJ, et al. (year?) Cohesin-dependent and independent mechanisms support chromosomal contacts between promoters and enhancers.

73. Kojic A, et al. (2018) Distinct roles of cohesin-SA1 and cohesin-SA2 in 3D chromosome organization. Nat. Struct. Mol. Biol. 25(6):496–504.

74. Li Y, et al. (2020) The structural basis for cohesin–CTCF-anchored loops. Nature 578(7795):472–476.

75. Lhoumaud P, et al. (2019) NSD2 overexpression drives clustered chromatin and transcriptional changes in a subset of insulated domains. Nat. Commun. 10(1):4843.

76. Luo Z, Rhie SK, Lay FD, Farnham PJ (2017) A prostate cancer risk element functions as a repressive loop that regulates HOXA13. Cell Rep. 21(6):1411–1417.

77. Belaghzal H, Dekker J, Gibcus JH (2017) Hi-C 2.0: An optimized Hi-C procedure for high-resolution genome-wide mapping of chromosome conformation. Methods 123:56–65.

78. Casa V, et al. (year?) Redundant and specific roles of cohesin STAG subunits in chromatin looping and transcription control.

79. Martinez-Soria N, et al. (2019) The oncogenic transcription factor RUNX1/ETO corrupts cell cycle regulation to drive leukemic transformation. Cancer Cell 35(4):705.

80. Assi SA, et al. (2019) Subtype-specific regulatory network rewiring in acute myeloid leukemia.

81. Richart L, et al. (year?) STAG2 loss-of-function affects short-range genomic contacts and modulates urothelial differentiation in bladder cancer cells.

82. Gibcus JH, et al. (2018) Mitotic chromosomes fold by condensin-dependent helical winding of chromatin loop arrays.

83. Niskanen H, et al. (2018) Endothelial cell differentiation is encompassed by changes in long range interactions between inactive chromatin regions. Nucleic Acids Res. 46(4):1724–1740.

84. Nair SJ, et al. (2019) Phase separation of ligand-activated enhancers licenses cooperative chromosomal enhancer assembly. Nat. Struct. Mol. Biol. 26(3):193–203.

85. Ray J, et al. (2019) Chromatin conformation remains stable upon extensive transcriptional changes driven by heat shock. Proc. Natl. Acad. Sci. U. S. A. 116(39):19431–19439.

86. Zhao Q, et al. (2020) Quantitative trait loci mapped for TCF21 binding, chromatin accessibility and chromosomal looping in coronary artery smooth muscle cells reveal molecular mechanisms of coronary disease loci.

87. Shi C, et al. (2021) Chromatin looping links target genes with genetic risk loci for dermatological traits. J. Invest. Dermatol. 141(8):1975–1984.

88. Ghurye J, et al. (2019) Integrating Hi-C links with assembly graphs for chromosome-scale assembly. PLoS Comput. Biol. 15(8):e1007273.

89. Morf J, et al. (2019) RNA proximity sequencing reveals the spatial organization of the transcriptome in the nucleus. Nat. Biotechnol. 37(7):793–802.

90. Barutcu AR, Blencowe BJ, Rinn JL (2019) Differential contribution of steady-state RNA and active transcription in chromatin organization. EMBO Rep. 20(10):e48068.

91. Akdemir KC, Others (2019) Chromatin folding domains disruptions by somatic genomic rearrangements in human cancers. Nat. Genet. 10.

92. Kantidze OL, et al. (2019) The anti-cancer drugs curaxins target spatial genome organization. Nat. Commun. 10(1):1441.

93. Canzio D, et al. (2019) Antisense lncRNA transcription mediates DNA demethylation to drive stochastic protocadherin *α* promoter choice. Cell 177(3):639–653.e15.

94. Elbatsh AMO, et al. (2019) Distinct roles for condensin’s two ATPase sites in chromosome condensation. Mol. Cell 76(5):724–737.e5.

95. Dixon JR, et al. (2017) An integrative framework for detecting structural variations in cancer genomes.

96. Lai B, et al. (2018) Trac-looping measures genome structure and chromatin accessibility. Nat. Methods 15(9):741–747.

97. Kadota M, et al. (2020) Multifaceted Hi-C benchmarking: what makes a difference in chromosome-scale genome scaffolding? Gigascience 9(1).

98. Jacobson EC, et al. (2020) Hi-C detects novel structural variants in HL-60 and HL-60/S4 cell lines. Genomics 112(1):151–162.

99. Chignon A, et al. (2021) Enhancer-associated aortic valve stenosis risk locus 1p21.2 alters NFATC2 binding site and promotes fibrogenesis.

100. Raviram R, et al. (2018) Analysis of 3D genomic interactions identifies candidate host genes that transposable elements potentially regulate. Genome Biol. 19(1):216.

101. Pan DZ, et al. (2018) Integration of human adipocyte chromosomal interactions with adipose gene expression prioritizes obesity-related genes from GWAS. Nat. Commun. 9(1):1512.

102. Lopez-Pajares V, et al. (2017) 464 dynamic and stable enhancer-promoter contacts regulate epidermal terminal differentiation. J. Invest. Dermatol. 137(5):S80.

103. Norrie JL, et al. (2019) Nucleome dynamics during retinal development. Neuron 104(3):512–528.e11.

104. Stadhouders R, et al. (2018) Transcription factors orchestrate dynamic interplay between genome topology and gene regulation during cell reprogramming. Nat. Genet. 50(2):238–249.

105. Kieffer-Kwon KR, et al. (2017) Myc regulates chromatin decompaction and nuclear architecture during B cell activation. Mol. Cell 67(4):566–578.e10.

106. Collins PL, et al. (2020) DNA double-strand breaks induce H2Ax phosphorylation domains in a contact-dependent manner. Nat. Commun. 11(1):3158.

107. Kim YH, et al. (2018) Rev-erb*α* dynamically modulates chromatin looping to control circadian gene transcription.

108. Chan WF, et al. (2021) Pre-mitotic genome re-organisation bookends the B cell differentiation process. Nat. Commun. 12(1):1344.

109. Johanson TM, et al. (2018) Transcription-factor-mediated supervision of global genome architecture maintains B cell identity. Nat. Immunol. 19(11):1257–1264.

110. Chen CCL, et al. (2020) Histone H3.3G34-Mutant interneuron progenitors co-opt PDGFRA for gliomagenesis. Cell 183(6):1617–1633.e22.

111. Wang Y, et al. (2019) Reprogramming of meiotic chromatin architecture during spermatogenesis. Mol. Cell 73(3):547–561.e6.

112. Kriz AJ, Colognori D, Sunwoo H, Nabet B, Lee JT (2021) Balancing cohesin eviction and retention prevents aberrant chromosomal interactions, polycomb-mediated repression, and x-inactivation.

113. Jiang Q, et al. (2020) G9a plays distinct roles in maintaining DNA methylation, retrotransposon silencing, and chromatin looping. Cell Rep. 33(4):108315.

114. Barutcu AR, et al. (2018) A TAD boundary is preserved upon deletion of the CTCF-rich firre locus.

115. Zhu Y, Denholtz M, Lu H, Murre C (2021) Calcium signaling instructs NIPBL recruitment at active enhancers and promoters via distinct mechanisms to reconstruct genome compartmentalization.

116. Kaaij LJT, Mohn F, van der Weide RH, de Wit E, Bühler M (2019) The ChAHP complex counteracts chromatin looping at CTCF sites that emerged from SINE expansions in mouse. Cell 178(6):1437–1451.e14.

117. Gnan S, et al. (year?) Nuclear organisation and replication timing are coupled through RIF1-PP1 interaction.

118. Siersbæk R, et al. (2017) Dynamic rewiring of Promoter-Anchored chromatin loops during adipocyte differentiation. Mol. Cell 66(3):420–435.e5.

119. Du Z, et al. (2020) Polycomb group proteins regulate chromatin architecture in mouse oocytes and early embryos.

120. Tan L, et al. (2021) Changes in genome architecture and transcriptional dynamics progress independently of sensory experience during post-natal brain development. Cell 184(3):741–758.e17.

121. Schwarzer W, et al. (2017) Two independent modes of chromatin organization revealed by cohesin removal. Nature 551(7678):51–56.

122. Miura H, et al. (2019) Single-cell DNA replication profiling identifies spatiotemporal developmental dynamics of chromosome organization.

123. Brandão HB, et al. (2018) A mechanism of Cohesin-Dependent loop extrusion organizes mammalian chromatin structure in the developing embryo.

124. Chatzidaki EE, et al. (2021) Ovulation suppression protects against chromosomal abnormalities in mouse eggs at advanced maternal age.

125. Silva MCC, et al. (2020) Wapl releases scc1-cohesin and regulates chromosome structure and segregation in mouse oocytes. J. Cell Biol. 219(4).

126. Simmons E, et al. (2021) Independence of chromatin conformation and gene regulation during drosophila dorsoventral patterning. Nat. Genet. 53(4):487–499.

127. AlHaj Abed J, et al. (2019) Highly structured homolog pairing reflects functional organization of the drosophila genome. Nat. Commun. 10(1):4485.

128. Cattoni DI, et al. (2017) Single-cell absolute contact probability detection reveals chromosomes are organized by multiple low-frequency yet specific interactions.

129. Li, et al. (2015) Widespread rearrangement of 3D chromatin organization underlies Polycomb-Mediated Stress-Induced silencing.

130. Gisselbrecht SS, et al. (2020) Transcriptional silencers in drosophila serve a dual role as transcriptional enhancers in alternate cellular contexts. Mol. Cell 77(2):324–337.e8.

131. Schauer T, et al. (2017) The drosophila dosage compensation complex activates target genes by chromosome looping within the active compartment.

132. Ramírez F, et al. (2018) High-resolution TADs reveal DNA sequences underlying genome organization in flies. Nat. Commun. 9(1):189.

133. Loubiere V, Papadopoulos GL, Szabo Q, Martinez AM, Cavalli G (2020) Widespread activation of developmental gene expression characterized by PRC1-dependent chromatin looping. Sci Adv 6(2):eaax4001.

134. Zenk F, et al. (2021) HP1 drives de novo 3D genome reorganization in early drosophila embryos. Nature 593(7858):289–293.

135. Szabo Q, et al. (2018) TADs are 3D structural units of higher-order chromosome organization in drosophila.

136. Cubeñas-Potts C, et al. (2017) Different enhancer classes in drosophila bind distinct architectural proteins and mediate unique chromatin interactions and 3D architecture. Nucleic Acids Res. 45(4):1714–1730.

137. Wang Q, Sun Q, Czajkowsky DM, Shao Z (2018) Sub-kb Hi-C in D. melanogaster reveals conserved characteristics of TADs between insect and mammalian cells. Nature Communications 9(1):1–8.

138. Chathoth KT, Zabet NR (2019) Chromatin architecture reorganization during neuronal cell differentiation in drosophila genome. Genome Res. 29(4):613–625.

139. Rowley MJ, et al. (2019) Condensin II counteracts cohesin and RNA polymerase II in the establishment of 3D chromatin organization. Cell Rep. 26(11):2890–2903.e3.

140. Erceg J, et al. (2019) The genome-wide multi-layered architecture of chromosome pairing in early drosophila embryos. Nat. Commun. 10(1):4486.

141. Gutierrez-Perez I, et al. (2019) Ecdysone-Induced 3D chromatin reorganization involves active enhancers bound by pipsqueak and polycomb. Cell Rep. 28(10):2715–2727.e5.

142. Ulianov SV, et al. (2021) Order and stochasticity in the folding of individual drosophila genomes. Nat. Commun. 12(1):41.

143. Ulianov, et al. (2018) Nuclear lamina maintains global spatial organization of chromatin in drosophila cultured cells in 2018 IEEE International Conference on Bioinformatics and Biomedicine (BIBM). Vol. 0, pp. 2493–2493.

144. Kribelbauer JF, et al. (2020) Context-Dependent gene regulation by homeodomain transcription factor complexes revealed by Shape-Readout deficient proteins. Mol. Cell 78(1):152–167.e11.

